# Survival of selected lactic acid bacteria isolated from pineapple waste in refrigerated pineapple juices

**DOI:** 10.1101/396838

**Authors:** Jannette Wen Fang Wu, Lidieth Uribe, Rodolfo WingChing-Jones, Jessie Usaga, Natalia Barboza

## Abstract

The aim of this research was to isolate and identify lactic acid bacteria (LAB) from pineapple waste. The survival in refrigerated pineapple juices, of a selected isolate with potential probiotic properties, was also studied. The 16S rRNA and *pheS* partial genes were used to identify LAB, multilocus sequence typing (MLST) genes were used in order to separate strains grouping with *Lactobacillus casei* and *L. paracasei* phylogenetically. Tests for survival at pH 2.0, resistance to lysozyme and tolerance to bile salts were used to screen the strains for potential probiotic properties. A *L. fermentum* isolate was used for the survival study. Three types of pineapple juice made from pulp, a blend of pulp and peel (80:20), and peel extract were inoculated to approximately 10^6^ CFU/mL with *L. fermentum* and stored at 4 °C for up to six weeks. The physicochemical composition of juices, including concentration of fermentable sugars and organic acids, total solids content, soluble solids content, titratable acidity and pH, was determined during the survival study. Two genera and five species were identified. Pineapple juices supported the survival of *L. fermentum* during refrigerated storage but the population of the bacteria decreased over time regardless of the juice type. Juice made from pulp was a more suitable vehicle for the survival of the selected LAB. Some of the juice physicochemical properties, including sugars and organic acids content, pH and titratable acidity, varied significantly (P<0.05) during storage. Further sensory studies are necessary to evaluate consumer acceptance of juices containing the selected isolate.

**IMPORTANCE:** Lactic acid bacteria (LAB), isolated from pineapple waste, were phylogenetically analyzed and characterized in regards to their tolerance to pH 2.0, lysozyme and bile salts; showing their potential as probiotic strains, if health benefits associated to their ingestion are eventually confirmed. Moreover, pineapple juice supported the survival of *Lactobacillus fermentum*, isolated from the same food matrix, during refrigerated storage at 4 °C. Among the three pineapple juices tested (pulp, pulp + peel and peel), *L. fermentum* survived better in juice made from pulp. However, significant variations were observed overtime in some of the physicochemical properties of the juices including sugars and organic acids content, pH and total titratable acidity.

## INTRODUCTION

Probiotics are live microorganisms which, when administered in adequate amounts, confer health benefits on the host (1). Them have been traditionally used for the development of functional dairy products such as fermented milk and yogurt (2). Beneficial effects associated with the ingestion of these microorganisms include maintenance of a healthy gut microbiota, improvement of the digestibility of foods, alleviation of lactose intolerance, prevention of diarrhea, control of blood cholesterol levels and hypertension, stimulation of the immune system and prevention of atopic allergies (3-5). However, consumption of milk-based beverages may be limited for certain consumers due to allergies, cholesterol diseases, dyslipidemia, and vegetarianism (6). Alternative and innovative food matrices, including fruit juices (7), are currently being investigated as substrates for novel non-dairy functional foods. Fruit juices are of particular interest as new carriers of probiotic lactic acid bacteria (LAB) due to their high concentrations of nutrients, particularly sugars, which may encourage probiotic growth. Moreover, juices do not contain starter cultures that compete with probiotics for nutrients during storage (8). The minimum concentration of live probiotic bacteria at the expiration date of a beverage should be around 7 logarithms of CFU/ml (4). Therefore, assessment of the survival of a potential probiotic strain in a selected beverage is critical in the development of new functional fruit-based drinks, even when previous studies have shown the potential of the studied microorganism as a probiotic in other food products.

Pineapple (*Ananas comosus*) ranks third in world tropical fruit production, preceded by banana and citrus (9). Costa Rica is one of the world’s largest producers of the cultivar ‘MD-2’ (10) and it is among the top five pineapple-producing countries (Thailand, Brazil, Philippines, Costa Rica and India, in descending order) (http://www.fao.org/statistics/). In Costa Rica, in addition to traditional pineapple juice made from pulp, innovative beverages are produced from pineapple peels and blends of fruit peel and pulp, products that in other latitudes may be considered an agro-industrial waste. Approximately 30 % of pineapple is turned into waste during a canning operation. Pyar et al. (11) investigated the potential of pineapple waste as growth medium for *Lactobacillus* species and found that bacterial growth on pineapple-waste culture medium was comparable to growth on Man Rogosa and Sharpe (MRS) agar. Thus, it might be expected that LAB isolated from pineapple waste may show an enhanced survival response in pineapple-derived beverages, and therefore the relevance of its isolation and identification is of particular interest. Moreover, limited information is available regarding the factors influencing probiotic survival in fruit juices (12, 13). Studying the effect of species/strain, juice composition and storage temperature and time on the survival of selected probiotic bacteria will enhance the existing knowledge related to this topic (4).

The aim of this research was to identify potential LAB strains isolated from pineapple waste and to study the survival of a selected strain in three different pineapple juices made from pulp, peels and a blend of both. The effect of juice composition on survival of the selected LAB and the changes in juice pH, total titratable acidity, soluble and total solids content, sugars content (glucose, fructose and sucrose) and organic acid concentration (citric, lactic and malic acid) during refrigerated storage of the beverages was also investigated.

## RESULTS AND DISCUSSION

### Bacterial strains: isolation, characterization and molecular identification

Twelve LAB morphotypes were isolated from pineapple waste, 16S rARN and *phe*S genes were amplified and sequenced (Table 1, Table S1). Two genera were identified: *Lactobacillus* spp. (11 strains) and *Weissella* spp. (one strain). The first one was the most common finding which is consistent with other reports of bacteria isolated from fruits and vegetables where *Lactobacillus* is characterized by its exceptional size and genetic diversity (14, 15), which have hindered its taxonomy (16). A clear cluster was observed comprising the costarican sequences and Genbank’s selected sequences of the 16S rRNA region and *pheS* gene from bacteria (www.ncbi.com) (Fig 1). Five different species were identified: *L. parafarraginis* (2), *L. fermentum* (2) and *W. ghanensis* (1); additionally, seven isolates were grouped with *L. casei* and *L. paracasei*. However, it was not possible to separate both species considering partial genome sequences of the 16s rRNA and *pheS* genes (Fig. 1).

**TABLE 1.**
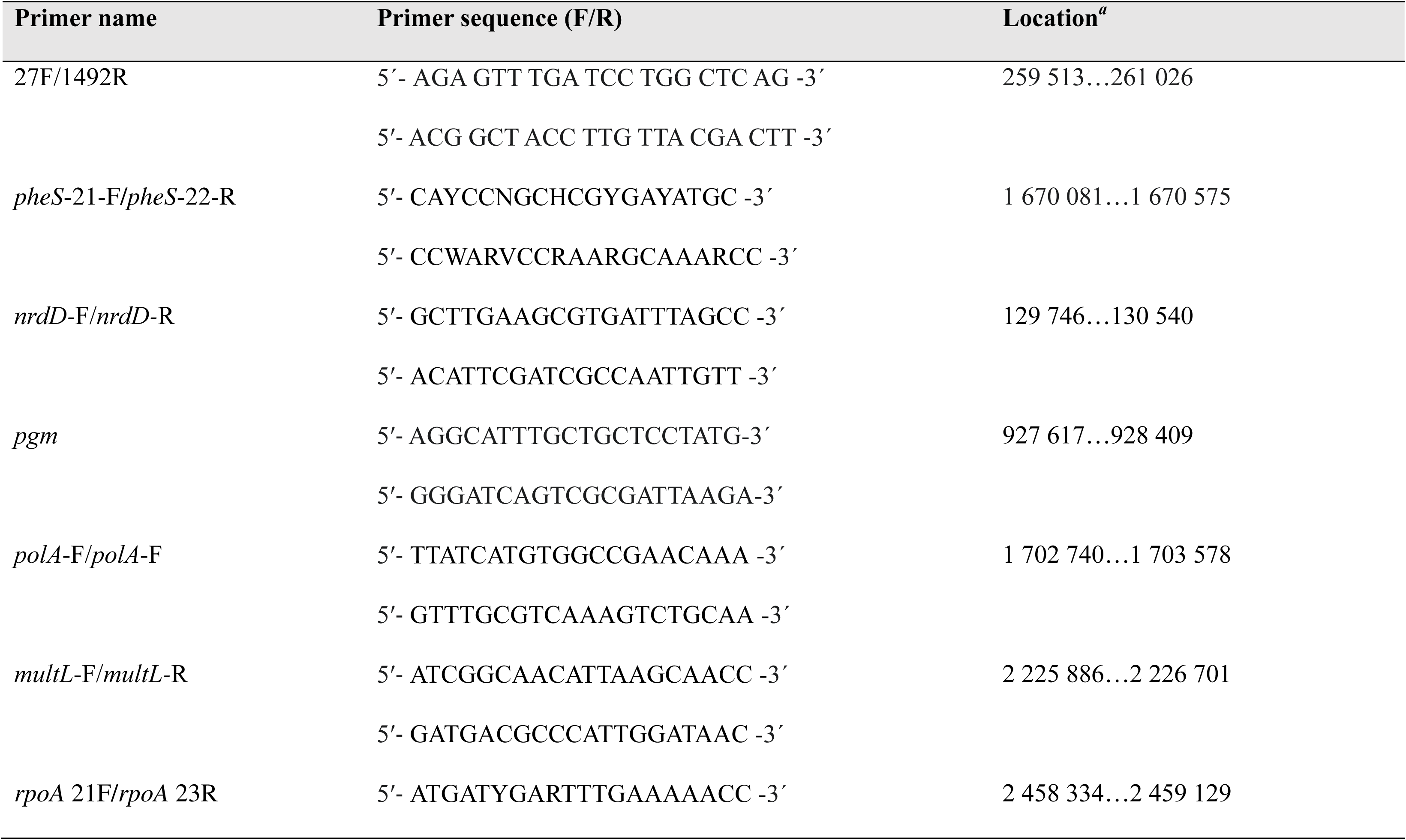
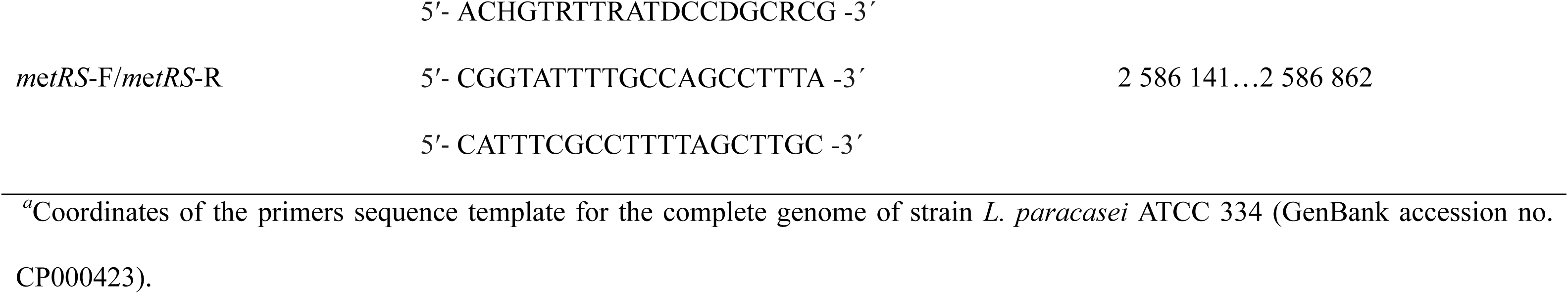
Primers used for molecular identification of lactic acid bacteria (LAB) isolated from pineapple waste residuals

**Fig 1.**
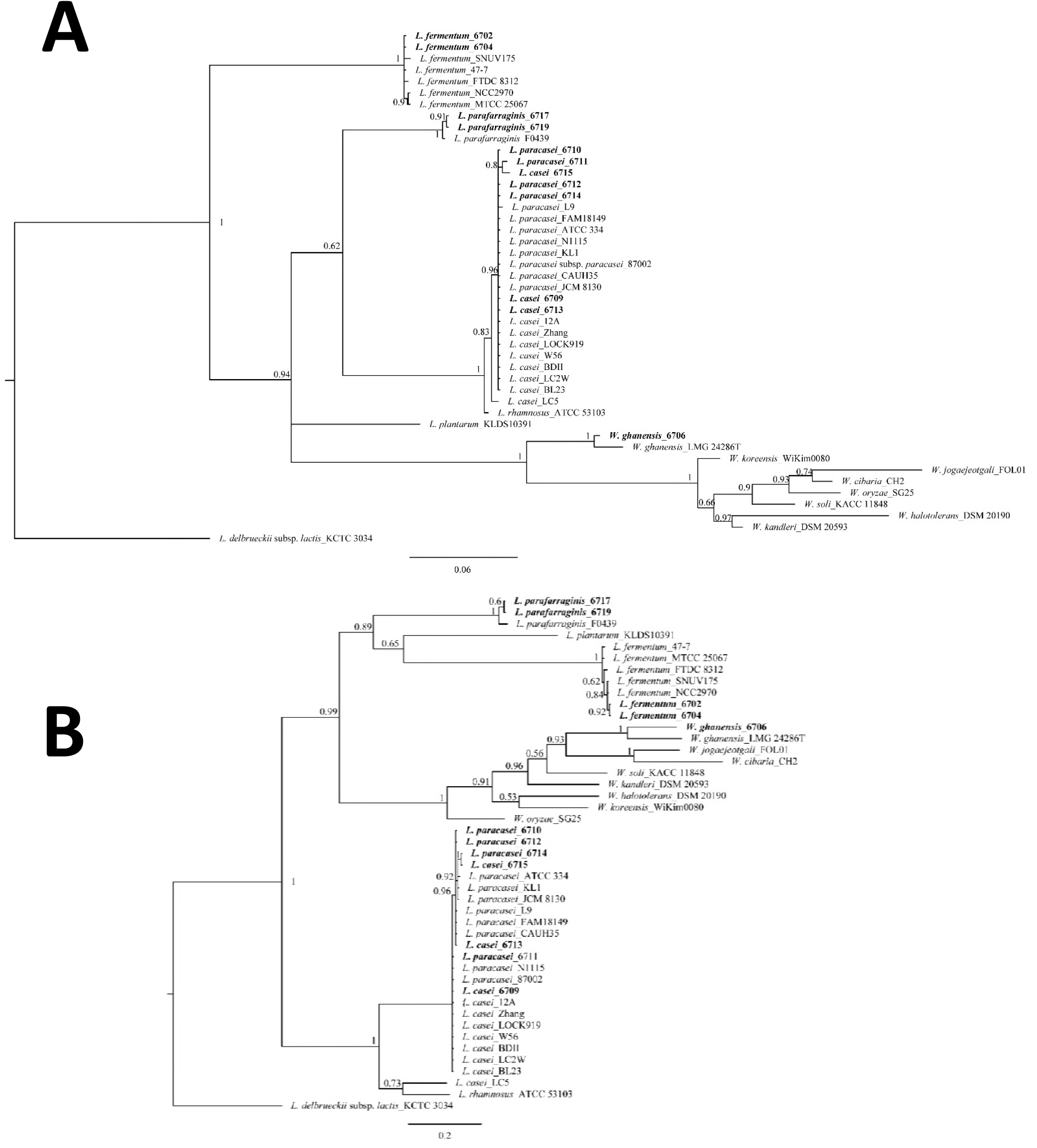
Phylogenetic tree for equivalent partial 16S rRNA gene sequences (1299 nucleotides (nt)) (A) and phenylalanyl-tRNA synthase (*phe*S) gene (420 nt) (B) of lactic acid bacteria’s (LAB) isolated from pineapple waste and comparison with sequences available in databases from other parts of the world. Sequences were aligned (using MUSCLE) and the phylogenetic tree was estimated by using Bayesian inference. Numbers above branches represent the Bayesian posterior probabilities. The bar indicates substitutions per site. Sequences corresponding to those present in Costa Rica are shown in bold letters. The phylogenetic trees were rooted using an equivalent sequence from *L. delbrueckeii* subsp. *lactis* KTCT 3034.

LAB strains have been isolated and characterized from different substrates in recent years because they potentially have probiotic properties and play an important role in the production of fermented food and beverages (17, 18). García et al. (19) for example, used the probiotic strain *L. fermentum* UCO-979C due to its potent antibacterial activity anti-*Helicobacter pylori.* Likewise, De Souza et al. (20) isolated *L. fermentum* and *L. casei* from mozzarella cheese in order to select novel and safe strains for the future development of functional fermented products; and Le and Yan (21) used probiotic strains of *Weissella* spp. to improve antioxidant activity during fermentation.

Equivalent length portions of the genes *nrdD, pgm, pheS, polA, mutL, rpoA* and *metRS* were analyzed and submitted to GenBank (Table S1) to resolve the species groups containing the isolated sequences of *L. casei* and *L. paracasei.* When compared to available sequences, a close relationship was found between our strains of *L. casei* and *L. paracasei*. Figure 2 depicts two groups of *L. casei* and *L. paracasei* strains, one containing *L. paracasei* (6710, 6711, 6712, 6714), *L. paracasei* L9 and *L. paracasei* ATCC 334 and the other comprising strains of *L. casei* and *L. paracasei* with no clear separation. It is important to mention that the taxonomic status and molecular identification of these species is still under debate (22 – 26) and is out of the scope of this paper, therefore, strains herein were named according to their closest match and phylogenetic position.

**Fig 2.**
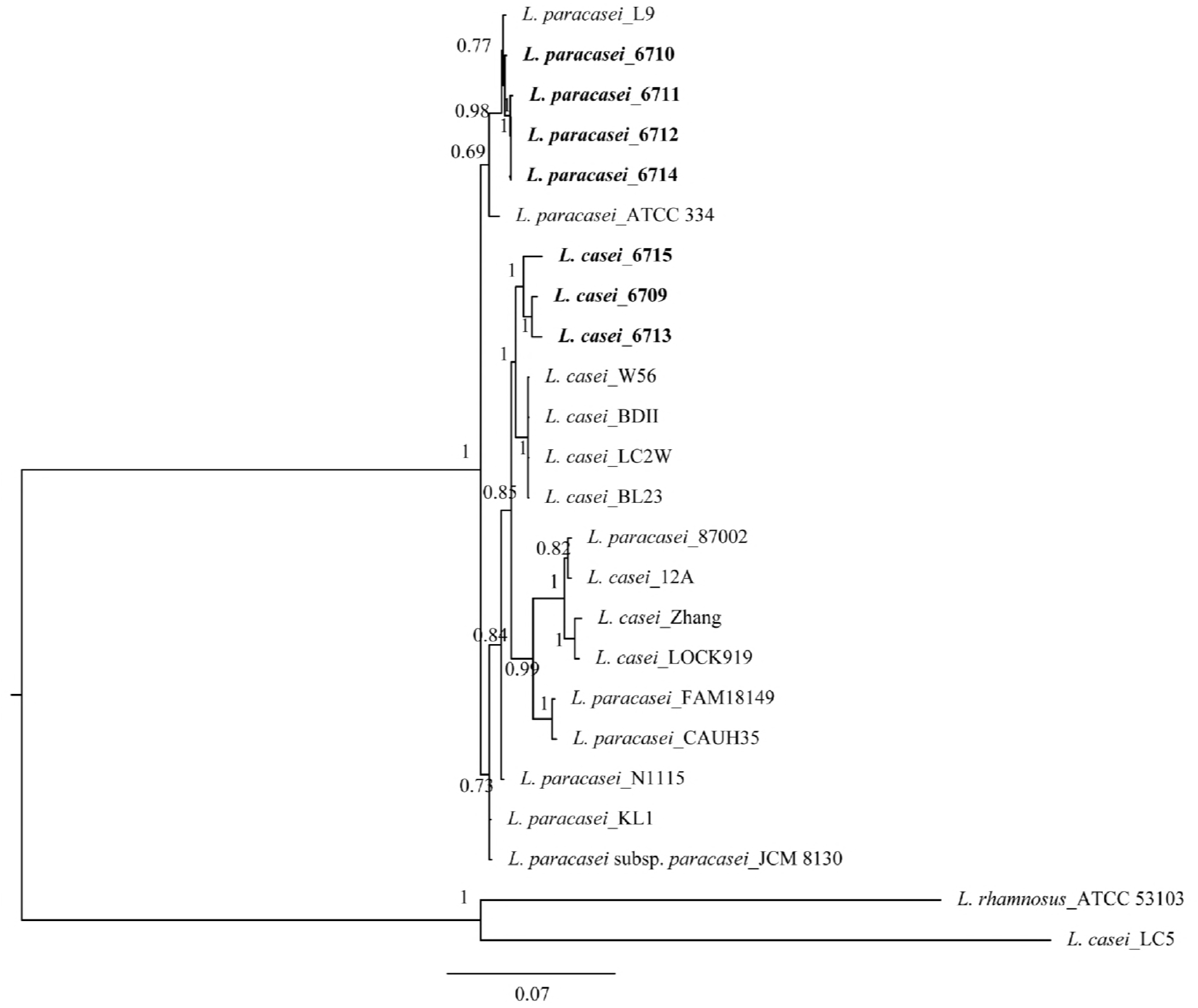
*L. casei* and *L. paracasei* phylogenetic tree considering the concatenated sequences of lactic acid bacterias (LAB) isolated from pineapple waste. Partial gene of anaerobic ribonucleoside-triphosphate reductase (*nrd*D) gene, DNA mismatch repair protein (*mut*L) gene, DNA polymerase I (*pol*A) gene, methionyl-tRNA syntethase (*met*RS) gene, phosphoglucomutase (*pgm*), polymerase alpha subunit (*rpo*A) and phenylalanyl-tRNA synthase (*phe*S) gene, were included. Comparison with sequences available in databases from other parts of the world was done. Sequences were aligned (using MUSCLE) and the phylogenetic tree was estimated by using Bayesian inference. Numbers above branches represent the Bayesian posterior probabilities. The bar indicates substitutions per site. Sequences corresponding to those present in Costa Rica are shown in bold letters. The phylogenetic trees were rooted using an equivalent sequence from *L. rhamnosus* ATCC 53103 and *L. casei* LC5.

### Tolerance to pH 2.0, lysozyme and bile salts

The tolerance to a pH of 2.0 of the 12 LAB isolates was evaluated. This condition simulates the phase of gastric emptying and is a strongly discriminative pH for the selection of high-acid tolerant strains (3, 27). All strains were viable after 3 h exposure to pH 2.0. However, populations were reduced (Table 2). As expected, the control samples (pH 6.0) did not show a log reduction during the time tested (data not shown). Survival was highest for *L. parafarragines* (6719) followed by *paracasei* (6710), *L. casei* (6715), and *L. fermentum* (isolates: 6702 and 6704). Overall, survival rates were greater than 80 % for most of the isolates, with the exception *L. casei* (6709) and *W. ghanensis* (6706). The high tolerance to a low pH shown by *Lactobacillus* gender was in concordance with previous reports (28), with exception of the isolate of *L. casei* (6709) which showed the lowest survival response. The tolerance in acid medium is one of the most important traits that could have the probiotic bacteria’s (29), because before they reach the gastrointestinal tract they must survive in the stomach. The average of pH during human digestion is around 3.0 to 2.0 with gradients from 4.0 to 1.8 during 2 h to 3 h period (30).

**TABLE 2.**
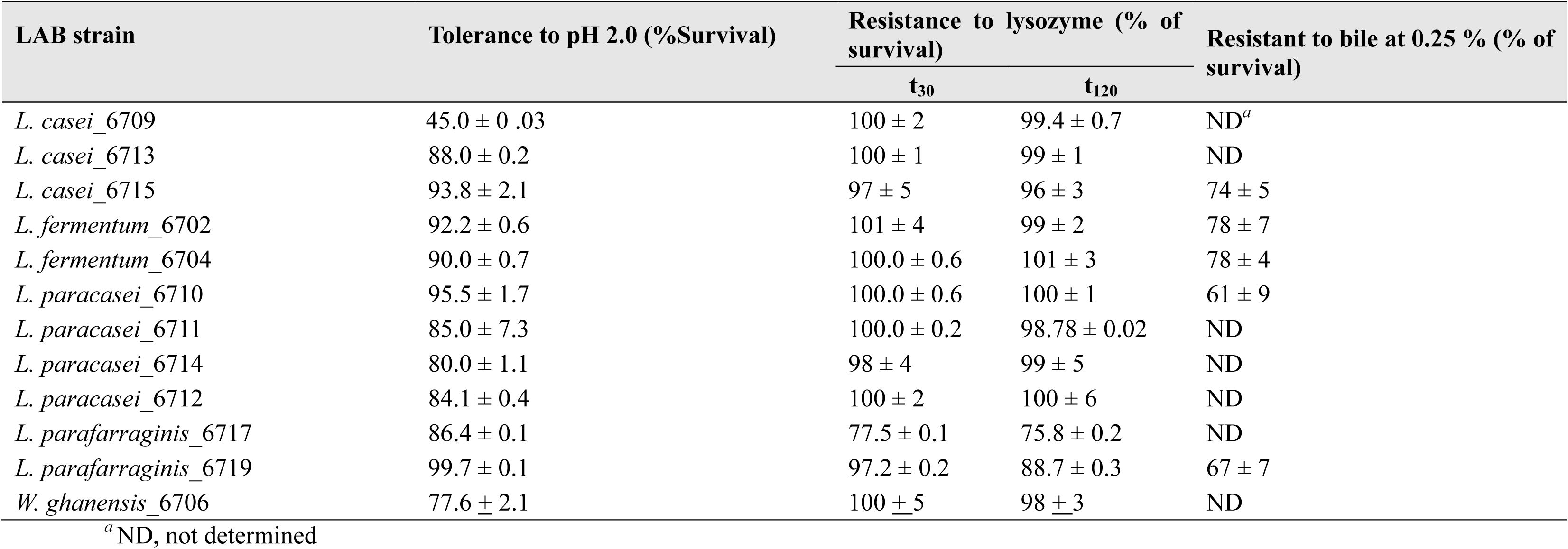
Tolerance to a pH of 2.0, lysozyme and bile salts of the LAB isolates (mean ± standard deviation, *n* = 3)

Survival after exposure to lysozyme for 30 min ranged from 77.5 to 100 % (Table 2). Similar values were found after 120 min of exposure. All LAB strains were also grown in medium without lysozyme and used as a control. These results confirmed the high resistance of most LAB isolates to lysozyme under conditions that simulate the *in vivo* dilution by saliva that occurs after juice ingestion. Bacterial resistance to lysozyme has been attributed to the peptidoglycan in the cell wall, the physiological state of the cell and the lysozyme in the medium (31). Similar results were found by García-Ruiz et al. (3), using LAB strains of *L. pentosaceus, L. casei* and *L. plantarum*, the authors found survival percentages > 80 % after 120 min of incubation, with exception of some strains with survival percentages around 50 %.

Five LAB isolates, with 90 % or higher survival at pH 2.0 and resistance to lysozyme, were selected and evaluated for resistance to bile salts. According to Dunne et al. (32), human bile concentrations range among 0.3 % to 0.5 %. For this reason, all the strains were evaluated on a medium with 0.25 % of bile showing survival values higher than 61 % for all strains. *L. casei* (6715) and both *L. fermentum* isolates showed higher values. Results are summarized in Table 2. The ability to survive in the presence of bile is an important characteristic of potential probiotic strains (3, 29). Mathara et al. (33) established bile tolerance limits for the selection of probiotic strains. Tolerance to 0.3 % bile salts with survival above 50 % was considered good. Our results were similar to ranges previously reported for *Bifidobacterium, Lactobacillus* strains, *P. pentosaceus* and certain yeasts (3, 34 - 37). High bile tolerance is desirable since its benefits colonization in the host gastrointestinal tract (38). In concordance with previously reports, our results showed that bile survivor was not related to LAB species but it was strain-specific (27, 39).

Based on results of the three previous essays, *L. fermentum* (6702) was selected for survival studies in pineapple juices due to its enhanced survival response. *Lactobacillus* strains are usually highly resistant to acids (40), and therefore they are commonly used as probiotics. *Lactobacillus fermentum, L. casei* and *L. paracasei* have been reported as probiotic species. Similarly, *Weissellea* spp. have been identified as probiotics but it has also been associated with undesirable food changes and fruit deterioration (15).

### Physicochemical characterization of pineapple juices during storage

The effect of storage time on the physicochemical characteristics of the inoculated juice was significant (P<0.05) for all variables except juice total solids content. These differences were significantly dependent (P<0.05) on the initial *L. fermentum* population of the inoculum. Means, coefficients of determination (R^2^) and probabilities are listed in Table 3. The effect of juice type on total titratable acidity was significant (P<0.0001). Differences in total titratable acidity of pulp juice and pulp + peel juice were not significant (P>0.05); however, the titratable acidity of both juices was significantly different (P<0.05) from that of juice extracted from pineapple peels.

**TABLE 3.**
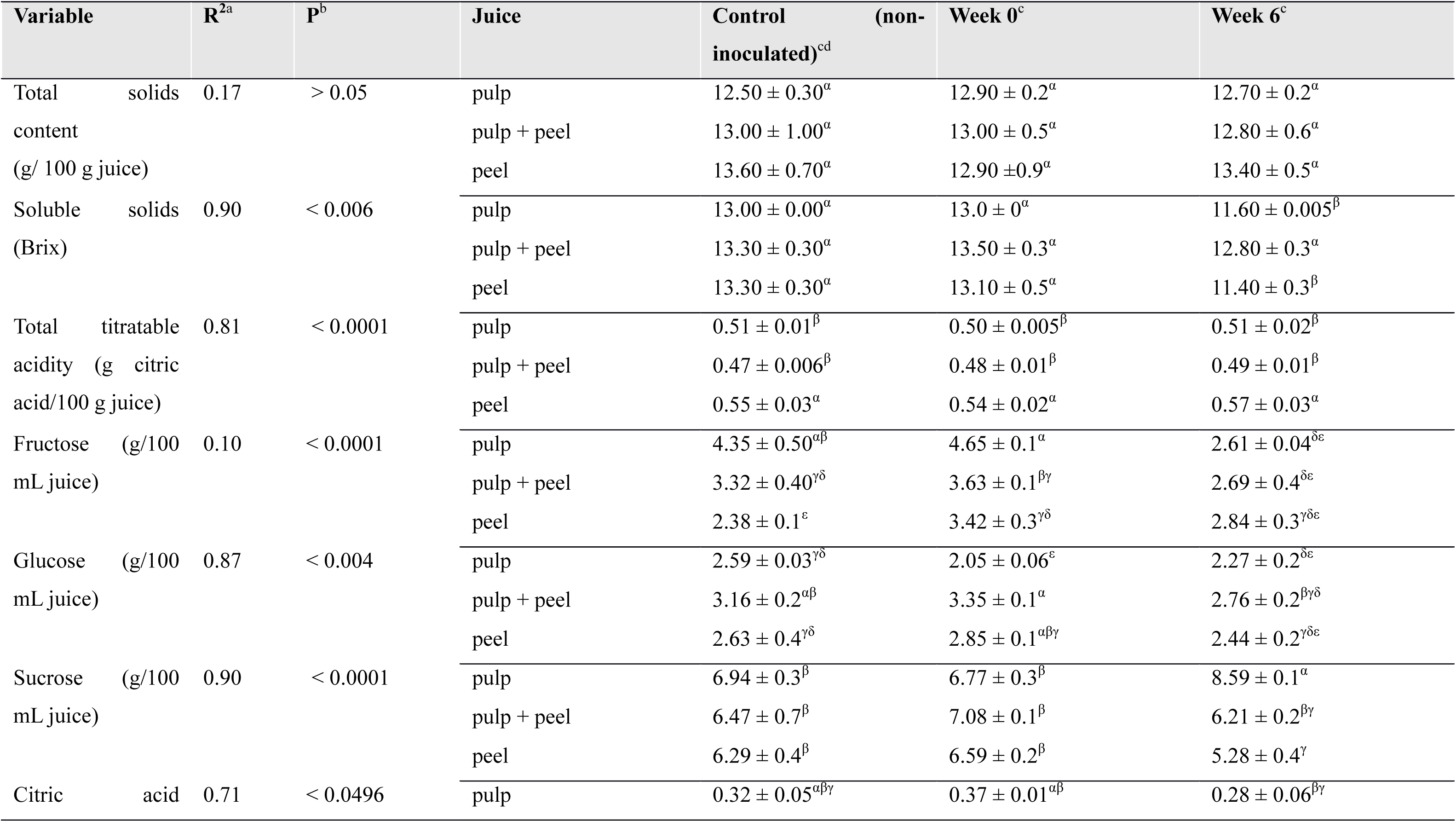
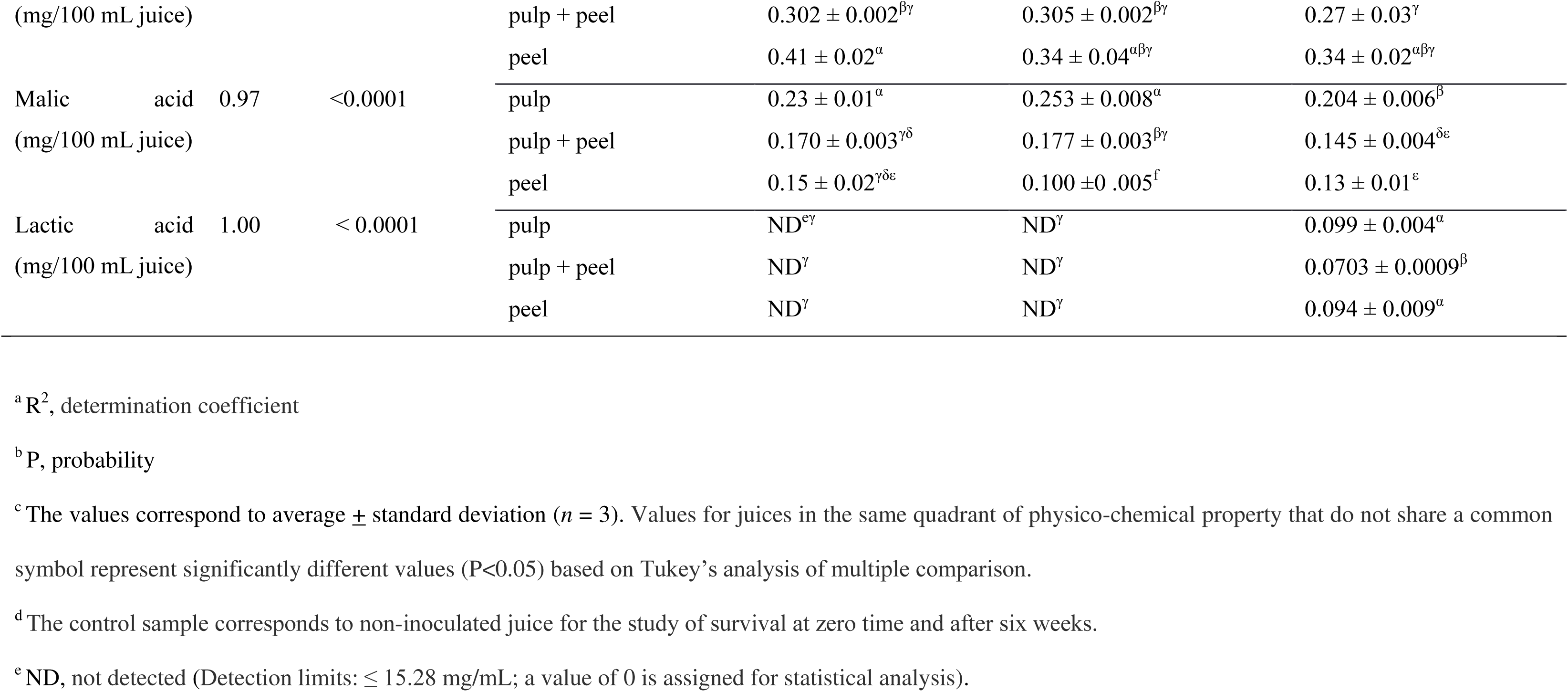
Physicochemical characterization of pineapple juices used to study *L. fermentum* survival at time zero and after six weeks of storage at 4 °C

For the case of the soluble solids content (° Brix), a significant interaction between storage time and type of juice was found (P = 0.006). After six weeks of storage, the soluble solids content of pulp and peel juices was lower than the initial Brix level. This result can be explained by the presence of the metabolically active *L. fermentum* strain inoculated into the samples.

The interaction between the effects of juice type and storage time on fermentable sugars was also significant, specifically, fructose (P<0.0001), glucose (P = 0.004) and sucrose (P<0.0001). Initial concentrations of the different sugars were consistent with previously reported data for pineapple juice; sucrose was the predominant sugar, followed by relatively equal fractions of fructose and glucose (41, 42). As observed in Table 3, a general downward trend in the sugar content after six weeks of storage was found. Initial and final fructose contents were significantly different (P<0.05). This difference was greater in juices made from pulp and pulp + peel. In the case of peel juice, no significant differences in fructose content were observed over time (P>0.05), but the glucose content differed significantly (P<0.05) after storage, possibly because of glucose metabolization by *L. fermentum*. For the case of sucrose, significant differences between the juices made from peel and pulp were observed.

The organic acids profile showed variations for the three acids analyzed (citric, malic and lactic). In all cases, significant interactions between juice type and storage time were found; citric (P = 0.0496), malic (P<0.0001) and lactic (P<0.0001). The three juices did not present significant differences of citric acid at week zero. At week six, the peel juice showed significant differences (P<0,05) when compared with juice made from pulp and pulp + peel. Citric and malic acid at week 6 were slightly reduced in comparison to the initial content. Lactic acid was detectable at week 6 in all samples. The reduction in citric acid content may be due to metabolization by the *L. fermentum* added to the juice (43). As with other members of the genus *Lactobacillus* spp., *L. fermentum* can metabolize malic acid through the formation of oxaloacetate and by the malolactic enzyme (44). As expected, an absence of lactic acid at week zero in all the juices was observed (41). Thus, the appearance of lactic acid at week six in all samples shows clear evidence of the metabolic activity of *L. fermentum* (45). Although the amounts produced were small, they were within the range previously reported (4) when the survival of *L. plantarum* in pineapple juice during refrigerated storage for up to six weeks was studied.

### *Lactobacillus fermentum* survival in pineapple juices

Survival curves for *L. fermentum* (storage at 4 °C) in the different pineapple juices and the evolution of juice pH during the experiment are shown in Fig. 3. Total solids and soluble solids, total titratable acidity and sugar and organic acid concentrations at the beginning and the end of the survival experiment are summarized in Table 3. The interaction between the nominal factors, juice type and storage time, was significant (P<0.0008). Thus, the effect of juice type on the survival of the bacteria was dependent on the storage time, and vice versa. A reduction in the initial populations of *L. fermentum* in the three pineapple juices was observed during storage (Fig. 3A). This effect was less pronounced in juice made from fruit pulp, even though this beverage had a lower pH than those containing peels (Fig. 3B). For juices containing peels, the cell concentration was lower than 10^7^ CFU/ml after one week of storage, in contrast to pulp juice in which the cell concentration decreased below 7 logs during the second week of refrigerated storage. Bacterial counts in juices containing pineapple peel extracts did not differ significantly at week six (P>0.05). Worth noting, the main metabolic substrates for lactic acid bacteria are fermentable sugars (fructose, glucose and sucrose); other carbon compounds such as organic acids can be used as secondary substrates (46, 47). This suggests that survival of *L. fermentum* is limited by the content of these carbon sources in the food matrix into which it is inoculated. Pulp juice showed the major content of fermentable sugar wich is coherent with the highest bacteria survival. Pulp + peel juice and peel juice showed the lowest fermentable sugars content, respectively. This results matched with a lower bacterial count and suggest the importance of sugar contain on bacterial survival. Therefore, the addition of fermentable sugars to the juice may be an option for reducing the inactivation rate of *L. fermentum* in pineapple-derived beverages. Furthermore, juice pH decreased slightly during the first two weeks before stabilizing. According to Ramos et al. (48), the presence of active and viable *L. fermentum* implies that its natural metabolic cycle is being completed. This leads to an increased production of organic acids, mainly lactic acid, causing the pH to decrease. Moreover, juice pH showed no further decrease after the second week of storage, which may be the result of the reduction of the bacteria in the juice over time (Fig. 3).

**Fig 3.**
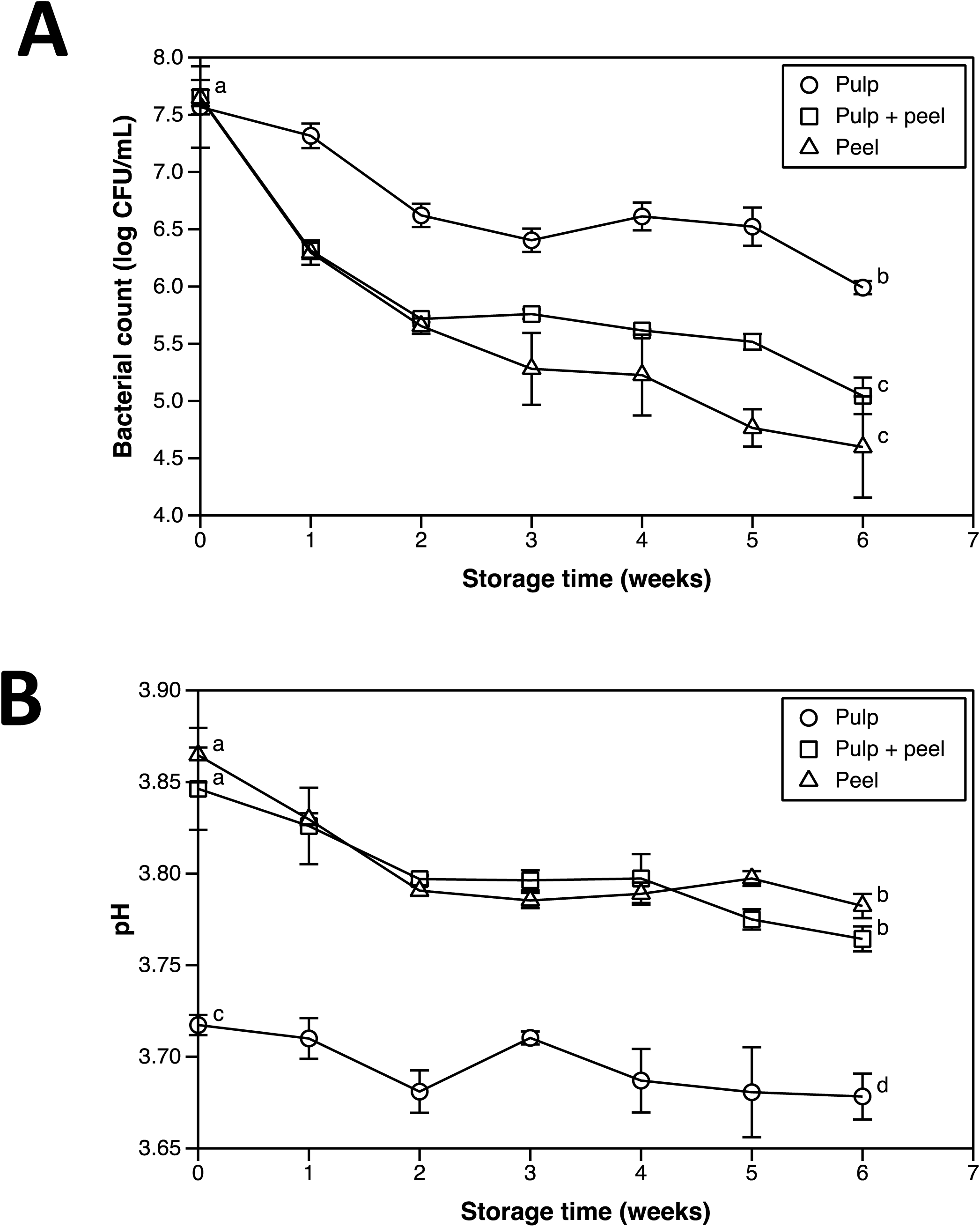
Evolution of the (a) *L. fermentum* population and (b) juice pH (mean ± standard deviation, *n* = 3) in pineapple juices (from pulp, peels and a blend of pulp and peels) during juice storage at 4 °C for up to six weeks. Values that do not share a common letter represent significantly different values (P<0.05) based on Tukey’s analysis of multiple comparisons.

Tests for tolerance to pH 2.0, resistance to lysozyme and tolerance to bile salts performed showed that lactic acid bacteria isolated from pineapple waste, specifically *L. fermentum* have potential as probiotic strains, if health benefits to the host are confirmed in further studies. Moreover, pineapple juice supported the survival of *Lactobacillus fermentum* during refrigerated storage. Among the three pineapple juices tested (pulp, pulp + peel and peel), *L. fermentum* survived better in juice made from pulp. However, regardless of the juice used, significant variations were found for some of the physicochemical properties of the juice, such as sugars and organic acids content, pH and titratable acidity. Further sensory studies are necessary to confirm the palatability of juices containing the selected isolate. Likewise, for a commercial application of this isolate, the study should be reproduced using frozen or freeze-dried cultures instead of the fresh cells used in our study.

## MATERIALS AND METHODS

### Isolation of bacterial strains

Twelve bacterial isolates obtained from pineapple waste from a local pineapple juice manufacturing company (San Jose, Costa Rica) were tentatively identified as LAB. Isolates were cultured on MRS agar (DIFCO, France) incubated at 35 ± 2 °C for 24 h in anaerobic conditions and subjected to Gram staining for morphology and preliminary identification. LAB cultures were maintained in 20 % (v/v) glycerol (a cryo-preservative that serves as a carrier for the microorganisms) and stored at −80 °C until used.

### DNA extraction and PCR amplification

DNA of LAB isolates was extracted using a miniprep protocol described by Birnboim and Doly (49). Part of 16S rRNA gene (∼1500 bp) using the primers 27F (50) and 1492R (51) was amplified and sequenced. The reaction was performed in a final volume of 25 μl and a final concentration of 1 X reaction buffer, 1.5 mM MgCl_2_, 0.2 μM of each primer, 0.2 mM dNTPs, 1 U Taq DNA polymerase (Bio-Rad, USA) and 50 ng of DNA. The PCR program consisted of an initial denaturation step at 94 °C for 1 min, followed by 30 cycles of 40 s at 94 °C, 55 °C for 1 min and 72 °C for 1 min. The final extension step was 5 min at 72 °C.

In order to confirm previously results, phenylalanyl-tRNA synthase (*phe*S) gene was sequenced for all the LAB isolates. The reaction was performed using primer combinations pheS-21-F/pheS-22-R (52), the PCR product was amplified using iProof High-Fidelity DNA polymerase (Bio-Rad, USA). Initial denaturation at 98 °C for 30 s, followed by 35 cycles of 98 °C for 30 s, 60 °C for 30 s, and 72 °C for 30 s; final extension at 72 °C for 10 min; and holding at 4 °C. A 25 ml reaction was prepared according to iProof High-Fidelity DNA polymerase and 50 ng of DNA. PCR products were visualized after electrophoresis in 1 % agarose gel (high resolution) by staining with GelRed (10 000 X) (Biotum, USA). LAB isolates identify in *L. casei* and *L. paracasei* group were analyzed considering six more housekeeping genes: anaerobic ribonucleoside-triphosphate reductase (*nrdD*), alpha-phosphoglucomutase (*pgm*), DNA polymerase I (*polA*), DNA mismatch repair protein (*mutL*), polymerase alpha subunit (*rpoA*) and methionyl-tRNA syntethase (*metRS*) (Table 1). PCR was performed under the same conditions that *pheS* gene. The PCR products were sent to Macrogen^®^ (South Korea) for sequencing in both orientations. General criteria of selection were the position of the genes on the chromosome, previously reported polymorphism, and the presence of one single copy of the gene (17).

### Molecular data analysis

The obtained sequences were assembled using the Staden program (53). Sequence similarity searches were performed using BlastN in order to compare the obtained sequences with genome complete sequences available in the databases (54). All sequences were aligned using MUSCLE (55) and deposited in the GenBank (Table S1). For phylogenetic comparison of 16S rRNA gene and *phe*S gene; 12 BAL isolates identify on this research and 33 representative sequences from all the complete genome LAB available in GenBank (www.ncbi) for this species were included. A common sequence length of 1299 nucleotides (nt) for 16S rRNA gene and 420 nt of *phe*S gene were considered. For the analysis of *L. casei* and *L. paracasei* group Multilocus Typing Sequences (MTLS) of seven genes were considered and concatenated following the order. *nrdD* (734 nt), *pgm* (697 nt), *pheS* (420 nt), *polA* (731 nt), *mutL* (746 nt), *rpoA* (685 nt), *metRS* (636 nt). Sequences of seven LAB obtained on this research grouping with *L. casei* and *L. paracasei* and 17 LAB sequences available in GenBank (www.ncbi) were included on the study. Bayesian phylogenetic analysis was conducted using MrBayes (56). Analyses considered 10 million generations using eight chains and sampling every 2 000 generations and mixed model (57). The sequence of *L. delbrueckeii* subsp. *lactis* KTCT 3034 was included as an outgroup for phylogenetical analysis of 16S rRNA gene and *phe*S gene, while for the analysis of *L. casei* and *L. paracasei* group *L. rhamnosus* ATCC 53103 and *L. casei* LC5 were included as outgroup.

### Tolerance to pH 2.0

In order to select an acid tolerant strain for further studies, the 12 LAB isolates were exposed to pH 2.0 following the procedure of Ramos et al. (48). Lactic acid bacteria cultivated in MRS broth (DIFCO, France) at 35 ± 2 °C for 24 h were centrifuged (5000 rpm for 5 min at 24 °C) and washed two times in 0.1 % w/v peptone water, pH 7.0. Cells were resuspended in peptone water to a concentration of about 10^6^ colonies forming units (CFU/mL), then centrifuged and re-suspended in MRS broth (DIFCO, France) with pH adjusted to 2.0 using 1N HCl and incubated for 3 h at 35 ± 2 °C. Samples (10 mL) obtained at time 0 and after incubation for 3 h were grown in MRS agar plates incubated in anaerobic jars for 72 h at 35 ± 2 °C.

### Resistance to lysozyme

The lysozyme resistance assay was performed using the method described by Zago et al. (58) with some modifications. One mililiter of LAB cells were cultivated in MRS broth (DIFCO, France) at 30 ± 2 °C for 24 h, centrifuged at 5000 rpm for 5 min at 24 °C, washed two times in phosphate buffer (0.1 M, water pH 7.0) and resuspended in 2 ml of Ringer solution (8.5 g/L NaCl, 0.4 g/L KCl, 0.34 g/L hydrated CaCl_2_) (Sigma Aldrich, USA). To simulate an *in vivo* spit test, bacterial suspensions (10^8^ CFU/mL) were inoculated into a sterile electrolyte solution (SES) (0.22 g/L CaCl2, 6.2 g/L NaCl, 2.2 g/L KCl, 1.2 g/L NaHCO_3_) in the presence of 100 mg/L of lysozyme (Sigma Aldrich, USA). Bacterial suspensions in SES without lysozyme were included as controls. The survival rate was calculated as the percentage of CFU/mL after 30 and 120 min relative to the CFU/mL at time zero. The microbial population was determined by cell counts in MRS agar. Assays were carried out in triplicate. Serial dilutions were done in 0.1 % sterile peptone water. Plates were incubated for 72 h at 35 °C.

### Tolerance to bile salts

Isolates with a survival rate of 90 % or higher after exposure to pH 2.0 were evaluated for tolerance to bile salts. The method described by García-Ruiz et al. (3) was utilized with minor modifications. Strains grown overnight were inoculated (2 % v/v) independently into MRS broth with 0.25 % bile salt (w/v) (Sigma-Aldrich, USA). Cultures were incubated in tilted tubes at 35 ± 2 °C for 24 h at 250 rpm in a rotatory benchtop incubated shaker (Lab Companion model SI-600R, Jeio Tech Company, South Korea). LAB plate counts in MRS were determined and compared to a control culture (sample without bile salts). Results were expressed as the percentage of growth (CFU/mL) compared to the control. The assay was conducted in triplicate and each sample was plated in duplicate.

### Pineapple juices

Aseptic pineapple (*Ananas comosus*, cultivar MD-2) juice extracted from fruit pulp and two pineapple juice concentrates, one made from pineapple peels and the other from a blend of pineapple peel and pulp extracts (80:20, respectively), were obtained locally (San Jose, Costa Rica). The concentrates were reconstituted with distilled water to about 12° Brix. Reconstituted juices were pasteurized and hot filled at 83 °C into clean glass containers to ensure commercial sterility. Containers were immediately sealed and inverted for at least 5 min to sterilize the headspace and metal lids. The hot fill step for the juices made from concentrate was performed to obtain shelf stable samples and therefore avoid the presence of unwanted microbiota that would interfere with quantification of the selected LAB during the survival study. To verify the absence of background microbiota, three replicate samples of each juice were plated on plate count agar (OXOID Thermo Scientific Fisher, UK) and acidified potato-dextrose agar (pH 3.5) (OXOID Thermo Scientific Fisher, UK) and incubated at 35 ± 2 °C for 24 h and at 25 ± 2 °C for 5 days, respectively, to determine total plate counts as well as mold and yeast counts.

### Physicochemical characterization of juices

Juice pH was measured with a 450 pH/ion Meter (Corning, USA). Total titratable acidity, expressed in grams of citric acid per 100 g of juice, was measured with a Titrando 836 System (Metrohm, Switzerland). Total solids content was determined by the vacuum oven method (59). The soluble solids content was measured using an Abbe refractometer model NAR-4T (ATAGO, Japan). The concentration of sugars (sucrose, glucose and fructose) and organic acids (citric, malic and lactic) was assessed by High Performance Liquid Chromatography (HPLC, Shimadzu, Japan). To determine sugar concentrations, standard solutions (Sigma-Aldrich, USA) and samples were injected onto the HPLC unit provided with a refractive index detector (RID) and a Zorbax Carbohydrate Analysis column (150 mm x 4.6 mm ID, 5 μm particle size; Agilent Technologies, USA). Samples were resolved using a blend of acetonitrile and purified water (75:25 v/v) at a 1.2 mL/min flow rate in an isocratic run for 10 min at 30 °C. For the assessment of the concentration of organic acids, the HPLC unit was coupled with a photodiode array (PDA, model SPF-20^a^, Shimadzu) detection system and a Hi-plex H column (300 mm x 7.8 mm, 8 μm particle size; Agilent Technologies, USA). A solution of 0.0045 M H_2_SO_4_ was used as the mobile phase at a 0.6 ml/min flow rate in an isocratic run for 20 min at 50 °C. A detection wavelength of 210 nm was used. All physicochemical analyses were performed in triplicate.

### *Lactobacillus fermentum* survival in pineapple juices

#### Culture preparation

*Lactobacillus fermentum* (6702) isolated from pineapple waste from the same juice manufacturing company where juices were obtained was grown in MRS (DIFCO, France) broth and incubated at 225 rpm in a rotatory benchtop incubated shaker (Lab Companion model SI-600R, Jeio Tech Company, South Korea) at 35 ± 2 °C for 24 h. LAB cultures were preserved in 1.5 ml cryovials containing 30 % glycerol and stored at −20 °C until used. The inoculum concentration was approximately 10^9^ CFU/ml. The exact initial counts were determined in triplicate after inoculating the juice samples used in the survival experiment.

#### Experimental design

Survival of *L. fermentum* (6702) in three pineapple juices made from pineapple pulp, peel extract, and a blend of pulp juice and peel extract was evaluated using the procedure reported by Nualkaekul and Charalampopoulos (4) with minor modifications. Samples of 100 mL of the juices were independently inoculated with 0.07 mL of the selected LAB targeting an initial concentration between 10^7^ and 10^8^ UFC/ml. The inoculated juices were stored at 4 °C for up to 6 weeks. Samples were collected weekly and analyzed for pH, soluble solids content (expressed in °Brix) and LAB counts. To determine the count of viable LAB cells, 0.1 mL of juice was spread onto MRS agar (DIFCO, France) and incubated in anaerobic jars at 35 ± 2 °C for 72 h before counting. The survival experiments were performed in triplicate for each condition, using independent inocula for each of the replicates. To verify the absence of background microbiota, three replicate control samples of each juice were plated on plate count agar and acidified (pH 3.5) potato-dextrose agar and incubated at 35 ± 2 °C for 24 h and at 25 ± 2 °C for 5 days, respectively, to determine total plate and mold and yeast counts. The sugar content (glucose, fructose and sucrose), organic acid concentration (malic, lactic and citric), total solids content and total titratable acidity (expressed as percentage of citric acid in 100 mL of juice) of the juices were determined at the beginning and the end of the experiment.

The effect of storage time at 4 °C on the physicochemical properties of pineapple juices inoculated with a selected strain of *L. fermentum* was evaluated. A full factorial design with three levels for type of juice (pulp, pulp + peel and peel) and two levels for storage time (week 0 and week 6) was constructed. The initial LAB count was included in the model as an independent variable. In addition, the effects of storage time and juice type on the survival of *L. fermentum* in pineapple juices refrigerated at 4 °C were examined. A full factorial design with three juice types (pulp, peel + pulp and peel) and seven storage times (from week 0 to week 6) was used. The population of this microorganism was used as the response.

#### Statistical analysis

To analyze the physicochemical properties of the juices and the survival of *L. fermentum* over time, two- and three-way analyses of variance (ANOVA), Tukey’s honestly significant difference, and Student’s t tests for means comparisons were performed using JMP version 11 (SAS Institute Inc., USA). Differences were considered significant at a *P* value of < 0.05.

## Acknowledgments

This study was supported by the Costa Rican Ministry of Science and Technology (MICITT) and the University of Costa Rica (UCR), Project FI-031B-14. The authors thank Henry Castro and Arturo Pacheco (undergraduate research assistants, UCR) for their technical assistance in the Food Microbiology Laboratory at CITA and also María Fernanda Miranda for his technical assistance in the Molecular Biology Laboratory at CIBCM.

